# The Argo: A 65,536 channel recording system for high density neural recording *in vivo*

**DOI:** 10.1101/2020.07.17.209403

**Authors:** Kunal Sahasrabuddhe, Aamir A. Khan, Aditya P. Singh, Tyler M. Stern, Yeena Ng, Aleksandar Tadić, Peter Orel, Chris LaReau, Daniel Pouzzner, Kurtis Nishimura, Kevin M. Boergens, Sashank Shivakumar, Matthew S. Hopper, Bryan Kerr, Mina-Elraheb S. Hanna, Robert J. Edgington, Ingrid McNamara, Devin Fell, Peng Gao, Amir Babaie-Fishani, Sampsa Veijalainen, Alexander V. Klekachev, Alison M. Stuckey, Bert Luyssaert, Takashi D. Y. Kozai, Chong Xie, Vikash Gilja, Bart Dierickx, Yifan Kong, Malgorzata Straka, Harbaljit S. Sohal, Matthew R. Angle

## Abstract

Here we demonstrate the Argo System, a massively parallel neural recording system based on platinum-iridium microwire electrode arrays bonded to a CMOS voltage amplifier array. The Argo system is the highest channel count in vivo neural recording system built to date, supporting simultaneous recording from 65,536 channels, sampled at over 32 kHz and 12-bit resolution. This system is designed for cortical recordings, compatible with both penetrating and surface microelectrodes. We have validated this system by recording spiking activity from 791 neurons in rats and cortical surface Local Field Potential (LFP) activity from over 30,000 channels in sheep. While currently adapted for head-fixed recording, the microwire-CMOS architecture is well suited for clinical translation. Thus, this demonstration helps pave the way for a future high data rate intracortical implant.

## Introduction

Motor and sensory information are represented in the brain by coordinated ensembles of neurons with topographic maps that can span several centimeters in large animals and humans. Decoding information from these representations necessarily requires recording from large numbers of individual neurons with high temporal fidelity (Stevenson and Kording, 2011). As a result, there has been a recent impetus in both experimental and translational neuroscience to record from more neurons (Jun *et al*., 2017; Musk and Neuralink, 2019; Stringer *et al*., 2019; Obaid *et al*., 2020).

Within the field of experimental neuroscience, there have been significant advances in both electrical and optical methods for recording from large populations of neurons, each having its respective advantages. Optical methods for recording neural activity have made major strides through the development of fluorescent proteins such as genetically encoded calcium indicators (e.g. GCaMP6/7(Chen *et al*., 2013; Dana *et al*., 2019)) and voltage sensitive fluorescent proteins (e.g. Archon (Piatkevich *et al*., 2018) or QuasAR (Hochbaum *et al*., 2014; Piatkevich *et al*., 2018)). These new fluorescent probes enable functional imaging experiments that can simultaneously record from as many as 10,000 neurons in vivo (Hochbaum *et al*., 2014; Piatkevich *et al*., 2018; Stringer *et al*., 2019). While these are powerful experimental tools, approaches based on fluorescent proteins face significant barriers in clinical translation and can only record from shallow regions of the brain without implantable optics. Further, the expression of exogenous fluorescent proteins requires modification of host cells, which has substantial safety and regulatory implications when applied to humans. Lastly, the scattering of light in the brain and thermal sensitivity of brain tissue creates significant engineering challenges for developing a practically implantable imaging system that can spatially resolve activity without overheating tissue (Kozai and Vazquez, 2015; Acker *et al*., 2016).

By contrast, electrical recording is well established as a tool for basic science and clinical research (Strumwasser, 1958). Intracortical electrode arrays are used for brain-computer interface applications (Hughes *et al*., no date; Hochberg *et al*., 2012; Flesher *et al*., 2016; Lubin, Strebe and Kuo, 2017; Pandarinath *et al*., 2017) and intraoperative recording (Truccolo *et al*., 2011; Misra *et al*., 2014; Cash and Hochberg, 2015). The state of the art in clinical neurophysiology, the Utah Array (Rousche and Normann, 1998; Hochberg *et al*., 2012), Blackrock Microsystems, USA), has enabled several notable applications in neural prosthetics (Willett *et al*., no date; Flesher *et al*., 2016; Pandarinath *et al*., 2017), despite having a limited electrode count (100 electrodes) and covering only a small 4 mm × 4 mm cortical area. Success with the Utah Array has motivated efforts to record from more neurons over larger areas of the brain. The main approach has been the insertion of multiple Utah arrays in a single patient (Hughes *et al*., no date; Flesher *et al*., 2016), but the low density of electrodes and lack of multiplexing in the Utah array make the technology difficult to scale to higher channel counts.

Many recent efforts to ‘scale up’ neurophysiology have focused on lithographically patterned, thin film-based probes. Most notable are those based on active CMOS probes (e.g. Neuropixel (Jun *et al*., 2017)) and flexible thin-films stacks (e.g. Polyimide + metal (Chung *et al*., 2019; Musk and Neuralink, 2019)). These high channel-count devices have enabled novel experimental paradigms with acute and semi-chronic recording, but they have yet to be demonstrated as robust for chronic implantation in large animals. Thin silicon probes are fragile, and polymer-substrate probes commonly suffer from cracking (Kozai *et al*., 2015) and delamination between insulation and electrode (Ceyssens and Puers, 2015; Čvančara *et al*., 2020).

One technological approach that is both highly scalable and promises more immediate clinical application is the use of microwire-CMOS arrays (Kollo *et al*., no date; Obaid *et al*., 2020). These devices use arrays of parallel microwire electrodes, which are connected to active CMOS electronics for readout and stimulation. Microwire electrodes consist of a conductive metal wire core insulated by a solution-resistant dielectric such as a polymer or ceramic. Microwires have been used consistently and reliably over the last 70 years to record extracellular action potentials from the brains of experimental animal models and humans (Nicolelis *et al*., 2003; Jackson and Fetz, 2007; McMahon *et al.*, 2014; Schwarz *et al*., 2014). They are highly robust and suitable for chronic applications (Nicolelis *et al*., 2003; Jackson and Fetz, 2007; McMahon *et al*., 2014; Schwarz *et al*., 2014) and translational models (Hubel, 1959; Hubel and Wiesel, 1959; Bartels *et* al., 2008; Misra *et al*., 2014). Importantly, recent work with carbon-fiber based probes suggests that the foreign body response to inserted microwires can be dramatically reduced by using wires with diameters less than 20 µm (Kozai *et al*., 2012; Guitchounts *et al*., 2013; Patel *et al*., 2015).

Traditionally, microwire technology has been less scalable than integrated silicon probes due to the difficulty of connecting large arrays to a large number of amplifiers (Nicolelis *et al*., 2003; Schwarz *et al*., 2014; Sohal *et al*., 2014). Recently, however, simultaneous, parallel bonding of microwire arrays to high density CMOS sensor arrays have been successfully demonstrated (Obaid *et al.*, 2020), paving the way for large format arrays of many thousands of wires that would not otherwise be achievable.

To date, demonstrations of the microwire-CMOS concept have involved adapting CMOS arrays that were not specifically designed for the application and have limited channel count due to restrictions in the array readout (Obaid *et al*., 2020). To fully realize the potential of microwire-CMOS technology, we have designed a custom CMOS application-specific integrated circuit (ASIC), electronics, firmware, and software, providing an end-to-end solution for large scale recording.

This “Argo system” represents a significant advance in total data throughput, simultaneously addressing up to 65,536 channels at 32 kHz and 12 bits of resolution. The CMOS readout array can be bonded to microwire electrode arrays of varying electrode length, count, and spacing, creating a highly versatile system applicable to different experimental models. It is designed for recording in a head-fixed, in vivo preparation. Full sensor readout produces a data rate of up to 26 Gbps, which can be simultaneously streamed directly to disk and viewed in real-time through a web browser-based digital oscilloscope interface. To validate our system, we performed initial recordings in the rat cortex with arrays of up to 1300 penetrating microwires, detecting action potentials from 791 single-units. To demonstrate the large channel count recording capability of the device, we performed surface recording of stimulus-evoked local field potentials from the sheep auditory cortex with a microwire-based electrocorticography array with over 30,000 channels.

## Methods

### System overview

The Argo system is designed to enable simultaneous data acquisition on up to 65,536 channels at sampling rates of 32 kHz (Khan *et al*., 2018). The system consists of an array of platinum-iridium (PtIr) microwire electrodes, a custom CMOS voltage-amplifier array designed to read and amplify neural signals (Kollo *et al*., no date; Obaid *et al*., 2020), electronics to process and packetize these signals, and a computer that runs the custom data acquisition software and user interface server (Figure 1). Individual components are described in detail in subsequent sections.

**Figure 1:**
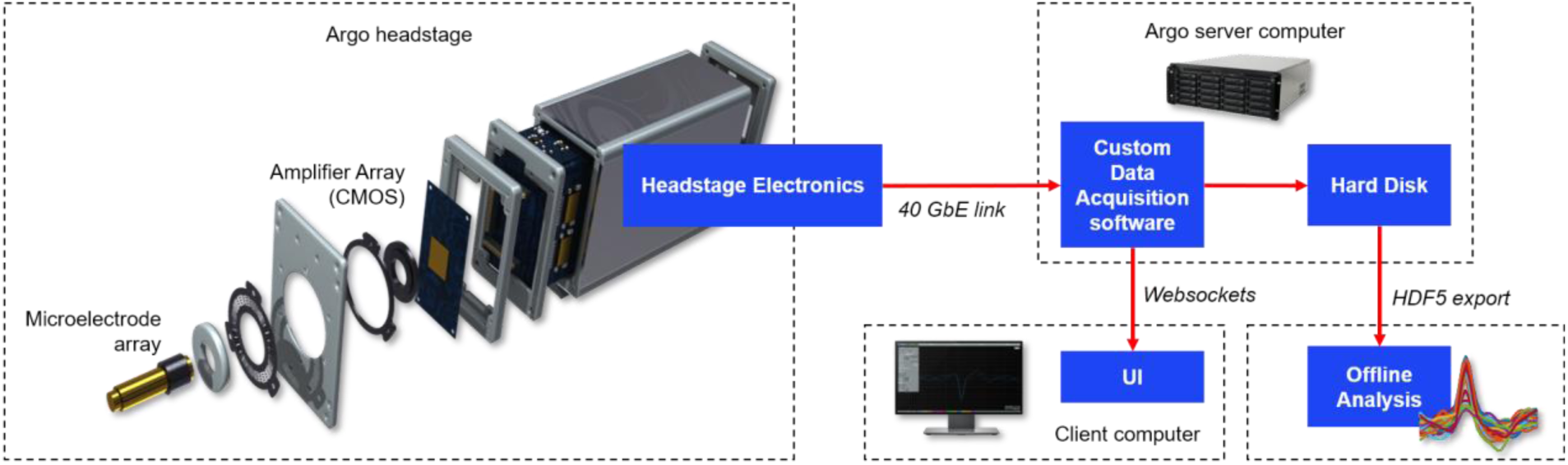
Block diagram of the Argo System. The Argo headstage comprises recording electrodes, the CMOS amplifier array, and electronics to digitize, packetize, and transmit signals over the optical data link. This data link connects it to the recording server computer, which hosts custom data acquisition software and a User Interface server. A client computer is used to read data from the server and display it in the system user interface. Data are exported as HDF5 files from the server and read into an offline processing computer where analysis and spike sorting are performed.

#### Microwire Arrays for Neural Recording

The recording array consists of a loosely ordered array of microwires (Tanaka Kikinzoku International, Japan) compressively and reversibly bonded to a custom CMOS amplifier array. The microwire cores are platinum-iridium alloy (90% Pt/10% Ir), selected for their biocompatibility and demonstrated recording/stimulation performance *in vivo* (Geddes and Roeder, 2003; Cogan, *2008*).

The fabrication process for our microwire arrays has been described previously (Obaid *et al.*, 2020). For intracortical arrays, the distal ends of the microwire electrodes are electro-sharpened in a parallel process adapted from (Musselman and Russell, 1990; Chang *et al*., 2012; Obaid *et al*., 2020). Microwires were bonded to a printed circuit board used to short all microwires and deliver electrical signals to them, and their distal ends were dipped in a 0.5 M CaCl_2_ solution. Sharp tips were formed by controlling the voltage applied to the wires in solution, and the speed at which wires were drawn out of solution. The amplitude, frequency, pulse duration, and duty cycle of the voltage applied to the wires were modulated using a Chroma Programmable AC Source (Model 61603, Chroma ATE, Taiwan). The wire speed was controlled using a stepper motor (ZST225B, ThorLabs, USA) and stepper motor controller (KST1, ThorLabs, USA), wires could be sharpened to arbitrary taper lengths and tip angles. The wire sharpening process consisted of four steps: (1) wire length equalization, (2) coarse electrosharpening, (3) fine electrosharpening, and (4) electrochemical polishing of the wire tips.(See Supplementary Table S1 for system parameters, and Supplementary Table S2 for sharpening process parameters). Between steps, the solution is circulated through a large bath using a preristaltic pump (Model 07528-10, Cole Parmer, USA) to evacuate reaction products. The final tip diameter is smaller than 200 nm (Figure 2A) to alleviate the “bed-of-nails”, or dimpling, effect that is commonly associated with the implantation of high density electrode arrays (<400 μm pitch) into cortical tissue (Obaid *et al*., no date; Nordhausen *et al*., 1992). Non-penetrating electrode arrays do not require this sharpening step, therefore tip preparation is performed later in the process.

**Figure 2:**
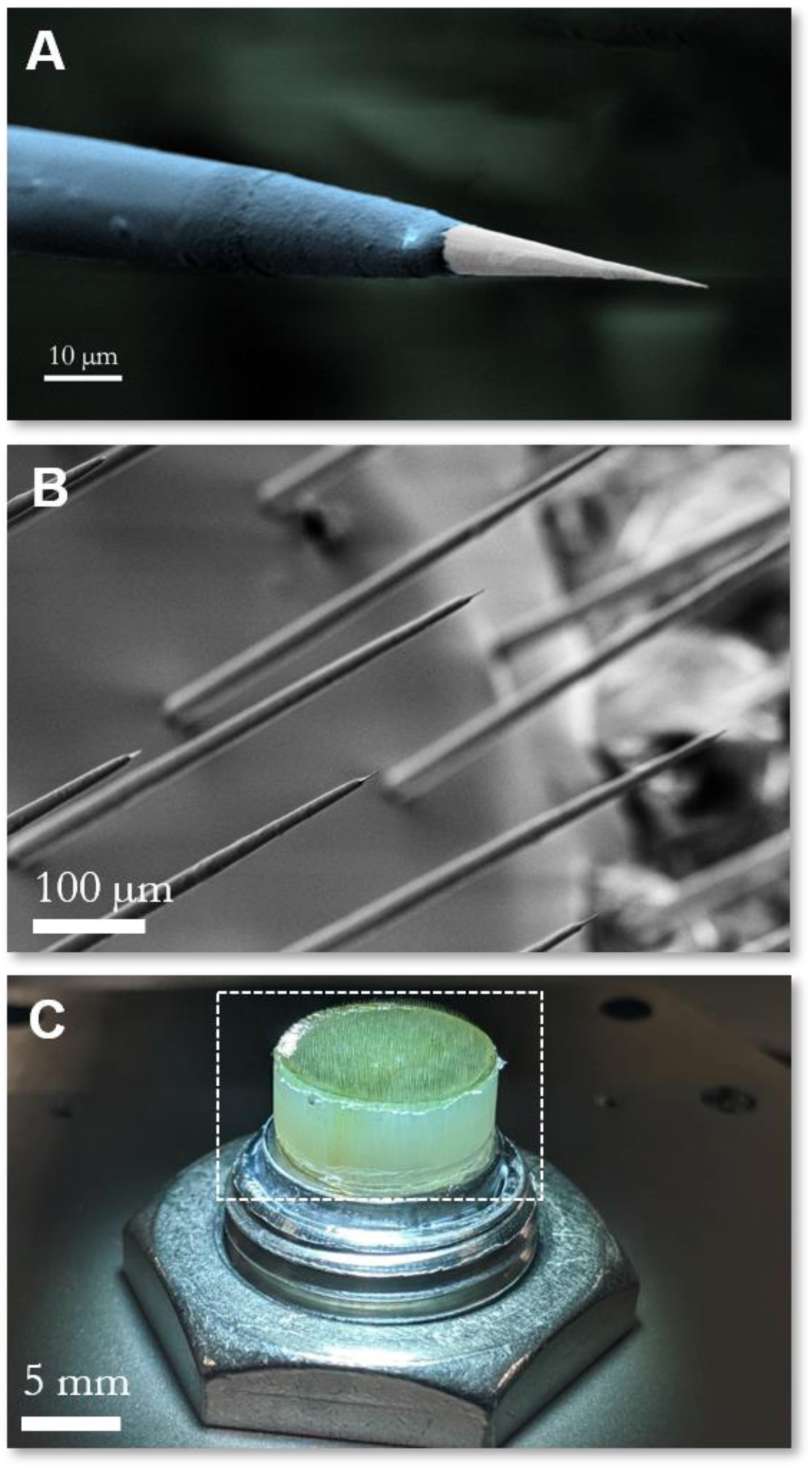
Microwire recording electrodes. (A) False color electron micrograph of a single electrosharpened microwire showing taper along the electrode length, alumina insulation layer (blue) and recording site at the tip (white), and (B) micrograph of the distal end of the microwire array showing a field of electrodes with electrosharpened tips. (C) Macroscopic optical image of the electrode array, highlighting the etched back recording end (white dashed box).

Microwires are then insulated with a thick sacrificial parylene-C layer to set the electrode spacing (100-400 μm). Following the coating step, the wires are tightly packed into a fluorinated ethylene propylene (FEP) heat shrink tube that is then shrunk to fix the wire positions within the array. Next, the proximal end of the array is prepared to ensure good electrical contact with the CMOS array when the array is mechanically pressed against the face of the sensor array (Obaid *et al*., 2020). This is achieved by polishing the proximal end of the array to be flat and then exposing the microwires at the proximal ends 10-20 μm by ashing in oxygen plasma using an SPI Plasma Prep III system (SPI Supplies, USA).

For non-penetrating electrode arrays, the tips are prepared by polishing the entire distal end of the array at this stage. This ensures that the recording sites are co-planar, smooth, and appropriately exposed.

For penetrating electrodes, the shank lengths of the distal ends (tips) of the bundled microwires are defined by ashing them with oxygen plasma as described above. At this stage, the arrays are coated with a robust 200-300 nm Atomic Layer-Deposited (ALD) alumina coating (Savannah S200, Cambridge NanoTech, MA, USA) to provide a high-quality insulation layer (Xie *et al*., 2014) that is then selectively de-insulated to define the length of the recording site at the wire tip.

Recording sites have a typical impedance of around 300-500 kΩ at 1 kHz in physiological saline. The smaller arrays used for *in vivo* experiments in rats had between 100 and 1300 recording electrodes, while the larger arrays prepared with flat tips for Local Field Potential (LFP) recordings in sheep had over 30,000 electrodes.

#### CMOS Sensor Design

The sensor in the Argo system is an application-specific integrated circuit (ASIC) designed to amplify and filter neural signals from a high-density microwire array. It was co-designed by Paradromics and Caeleste, CVBA (Belgium) and fabricated by X-FAB Silicon Foundries (Germany and Malaysia) in a 180 nm CMOS process technology node. The sensor consists of a pixel array of 256 × 256 pixels with a pixel pitch and dimensions of 50 µm x 50 µm, adding up to an active area of 12.8 mm x 12.8 mm for the readout array. The peripheral circuit elements for control and readout increase the total dimensions of the ASIC to 14.5 mm x 16 mm.

Each pixel has a 40 µm x 40 µm top metal pad which is used as the landing pad for the microwire electrode. This top metal pad is AC-coupled to a low-noise amplifier (LNA) chain. The LNA chain is composed of three main blocks: input amplifiers, an antialiasing low-pass filter, and an output column buffer (Figure 3).

**Figure 3:**
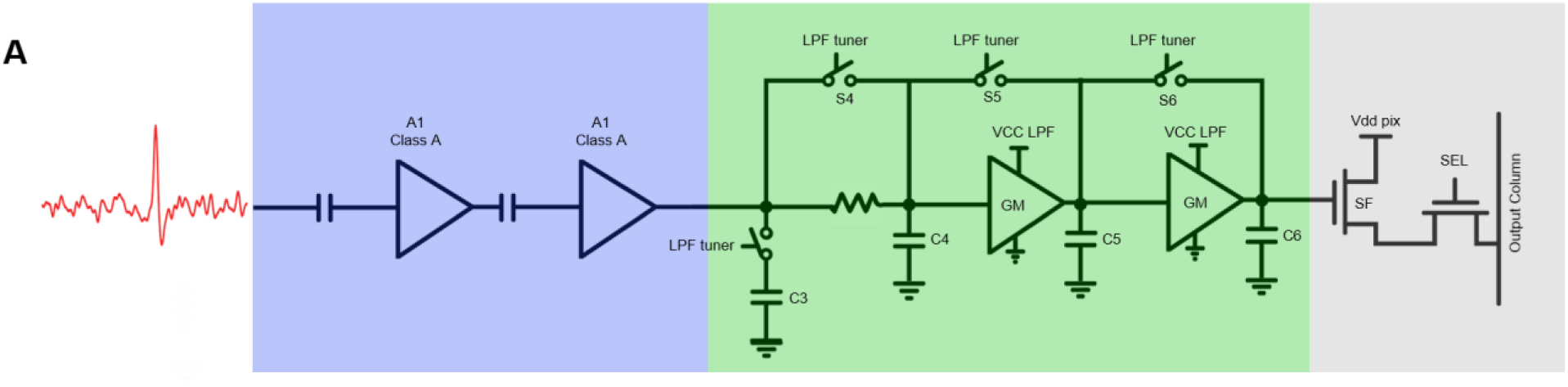
CMOS sensor characterization. The input signal is AC-coupled into two source followers biased in the Class A configuration (A1, A2, shown in blue), which together form the front-end low noise amplifier (LNA) chain. Each stage has a gain of 10 V/V. The third stage (green) is a third-order tunable low-pass filter that serves as an anti-aliasing filter. The final stages (grey) are for pixel selection to read out the stored value.

The amplifiers are implemented as common-source gain stages that are biased to operate in the class A regime. This topology was chosen due to its good linear response, high input impedance, and low noise. The drawbacks are higher power consumption and lower gain. To limit the power consumption and at the same time achieve the required gain of 100 V/V, the design implements two amplifiers in series, with each amplifier contributing a gain of approximately 10 V/V. In addition, each amplifier has a tunable input biasing circuit which in conjunction with an AC coupling input capacitor forms a tunable high-pass filter. Thus, the amplifier block provides a cascaded second order high-pass filter that has its 3dB point tunable from approximately 18 Hz to 300 Hz. This serves two functions. The first is the removal of DC offsets, drifts, and slowly varying out-of-band high amplitude signals. These undesired signal components can saturate the input and diminish the signal dynamic range, ultimately reducing the ability of the system to acquire meaningful signals. The second is to provide the flexibility of the system to operate in two modes. In one mode, setting the corner frequency to 18 Hz enables both neuron spiking activity and LFP to be recorded, since the latter has most of its integrated power below 100 Hz. In the second mode, LFP signals can be rejected by increasing the corner to higher frequencies. This allows for spiking signals to retain the entire dynamic range. Furthermore, the input impedance is sufficiently high as to not form a significant voltage divider with the otherwise high electrode impedance, which can range as high as 10 MΩ at 1 KHz. Specifically, the input impedance at 1 kHz has a resistive component of 4.4 GΩ and a capacitive component of 0.4 GΩ (400 fF).

To keep the integrated noise floor as low as possible while simultaneously providing antialiasing, the amplifier chain is followed by a third order low-pass filter with a tunable corner frequency between 8 kHz and 50 kHz. In our application all three corners are set to 20 kHz resulting in a 3dB point at approximately 12 kHz and providing good signal suppression at the Nyquist frequency of 16 kHz.

The output buffer of the pixel provides isolation between the pixel output and the column line used to multiplex the pixels in a single column. That is, when one of the pixels in the column is being read out the other pixels have their outputs disconnected from the line to avoid overloading. This reduces crosstalk between pixels and ensures that each pixel can read out signals from unique electrodes.

In the ASIC periphery, a set of control circuits and high-bandwidth amplifiers convert the multiplexed signal from single-ended to differential and bring the total gain of the signal chain to around 800 V/V. The sensor multiplexes 2048 individual channels to each of 32 high-speed analog outputs from the entire array. Furthermore, the output buffers are designed to drive long transmission lines leading to the external analog-to-digital converters ADCs.

#### Argo System Electronics

The system electronics consist of two custom printed circuit boards (PCBs). The first PCB (front-end board) is designed to house the CMOS sensor and support electronics. The second PCB (main board) is designed to digitize analog signals and deliver them to the server (Figure 4). It is designed in a rigid-flex form factor for the reasons of signal integrity, compactness, and ease of assembly.

**Figure 4:**
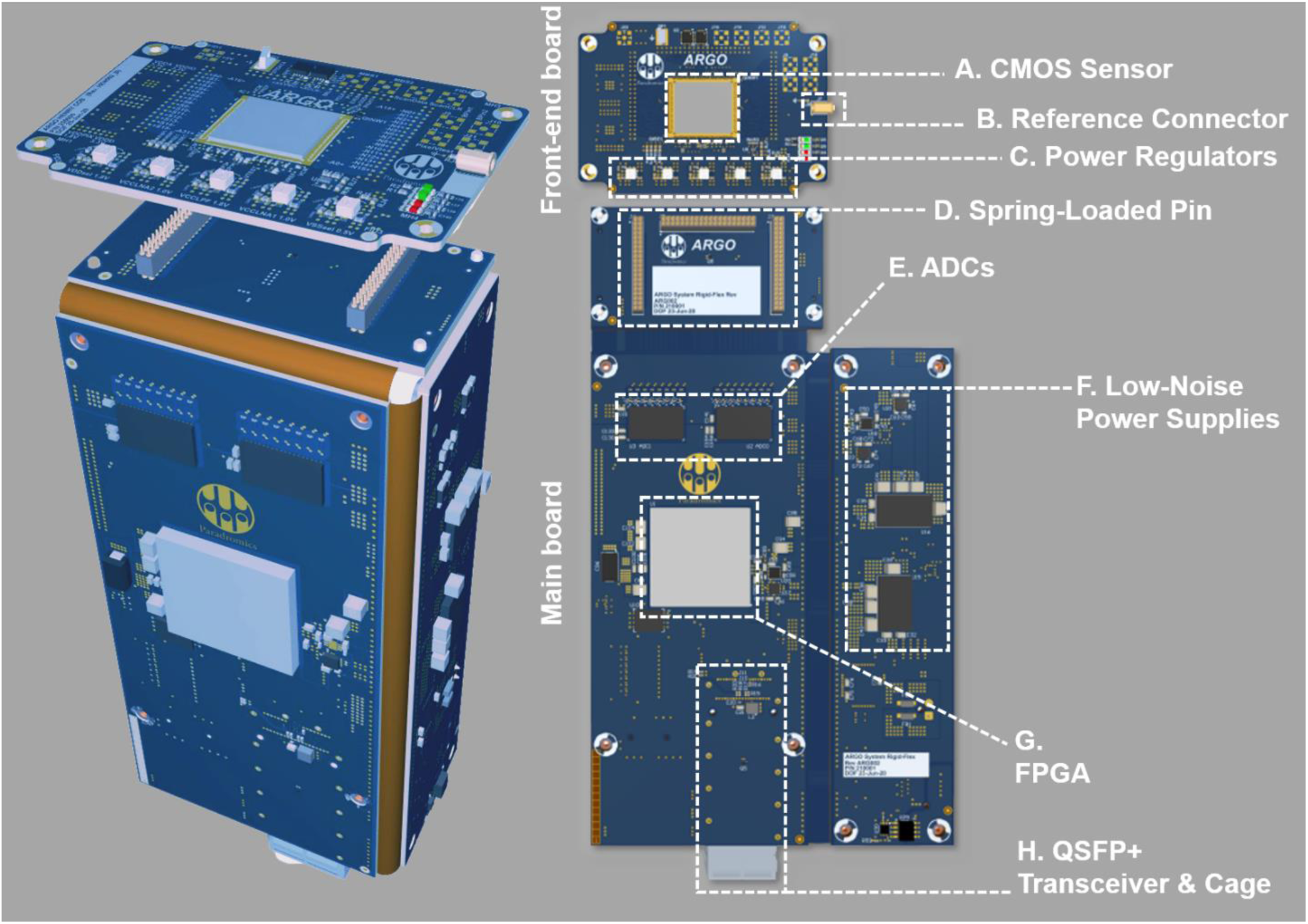
System electronics. The Argo system’s electronics are housed on two custom PCBs. (Left) PCBs are folded and aligned to fit the metal housing of the Argo system. (Right) A smaller PCB (the front-end board) holds the CMOS sensor (A), reference connector (B), and power regulators (C) for the sensor. This front-end board is connected to a larger main board via three banks of spring-loaded connectors (D). On this main board sit the analog-to-digital converters (ADCs, E), low-noise power supplies (F), the FPGA (G), and the Quad Small Form-factor Pluggable (QSFP+) transceiver and connector (H) over which data are transmitted using an optical fiber (not pictured).

The CMOS sensor is wire-bonded to the front-end board, which also contains a reference connector and a set of precision voltage regulators for the sensor. The front-end board is mounted in a metal housing designed to reliably connect with the microwire array after pressing (see Methods: Array connectivity testing for more details) (Kollo *et al*., no date).

The main board contains two 16-channel high-speed ADCs (ADS52J90, Texas Instruments, USA) as well as a Field-Programmable Gate Array (FPGA, XC7K160T-2FFG676-2, Xilinx, USA). The headstage requires an external 5V/3A DC power source and is kept galvanically isolated from powerline and other electronic noise sources, which can otherwise corrupt neural signals. The main board is housed within a separate headstage body designed to connect with the front-end main board frame. Electrical connections between the two boards are made via high-density spring-loaded pins, shown in Figure 4(D)) (855-22-040-30-001101, Mill-Max, USA). This allows reliable and repeatable mating between the boards without imposing extreme tolerance constraints on the fabrication of either the boards or the mechanical components. Two banks of pins transmit the multiplexed, high-speed analog output signals from the CMOS sensor to the ADCs (Figure 4(E)), while the third separate bank of pins transmits control signals from the FPGA to the CMOS sensor.

The ADC’s input signal range of 2 Vpp translates to an input signal range of ∼2.5 mVpp in the CMOS amplifiers, given the 800 V/V gain (see Methods: CMOS Sensor Design). The ADCs operate at a sampling frequency of 78.125 MHz, corresponding to a sensor sampling rate of 32 kHz, and digitize the analog signal to a 12-bit digital signal that is then transmitted to the FPGA over tightly impedance-controlled Low-Voltage Differential Signalling (LVDS) pairs. This high sampling rate provides further protection from aliasing artifacts. It also allows for spike sorting applications of the neural data, wherein a higher sampling rate leads to better cluster/unit separability in PCA space for an individual spiking channel (Lewicki, 1998; Ghanbari, Papamichalis and Spence, 2009).

The FPGA (Figure 4(G)) then demultiplexes the wide-band digitized signals from the 32 LVDS channels into the original 65,536 channels and remaps them to construct a 256 × 256pixel raster at 32 kHz. The logic in the FPGA generates timing signals for raster scanning the 256×256 pixels CMOS sensor array at a rate of up to 32,000 frames per second.

The FPGA is also used to coordinate data transfer to the host computer via a 40 gigabit Ethernet link through an OM3 optical fiber (943-99684-10005, Amphenol, USA). As the physical layer of the 40GBASE-SR4 protocol, the FPGA drives a Quad Small Form-factor Pluggable (QSFP+) optical transceiver (Figure 4(H)) through its 8 GTX transceivers. An embedded application processor (Xilinx Microblaze) within the FPGA fabric coordinates the ARP, ICMP, and UDP communication for control signals to and from the host computer. In order to maintain throughput for the high-speed neural data, the FPGA fabric directly assembles UDP packets using header information set by the embedded processor.

#### Computer systems for neural readout and data storage

To handle the large amount of data generated by the Argo system, the host computer is custom-built on a Supermicro dual Xeon server platform (X10DAC, Supermicro, USA) running Ubuntu Linux 18.04 Server Edition and equipped with two Intel Xeon E5-2640 v4 processors, 64 GB of ECC RAM and an Intel XL710 network interface card for 40GBASE-SR4 communication with the Argo headstage. This host server runs a fully custom data acquisition software package. This software produces two data streams: one raw data stream to be written to disk, and another stream to be served externally through a websockets interface, where data can be accessed through the application programming interface (API). The data acquisition software package is built on Intel’s Data Plane Development Kit (DPDK) to ingest the raw ethernet packets at full 40 Gbps line rate. The packets are evaluated for checksum integrity at the application level and are rearranged to be written to an array of 20 hard drives using Direct Memory Access (DMA) and a custom multi-threaded sharding system.

High-performance computing technologies such as core pinning, cache line optimization, and memory alignment are used throughout the data acquisition software stack to enable recording of full frame neural data at full 32 kHz frame rate. This allows the user to record data continuously for up to 8 hours, the duration of a typical acute sheep experiment, without running out of disk space or needing multiple recording systems in the surgical environment.

#### User interface for live data visualization

A custom UI for real-time monitoring was developed for the Argo system to facilitate in-experiment data validation required for collection of high-quality datasets. Some salient features of the UI include customizable bandpass filters (e.g. 300-6000 Hz for recording in the spike band), powerline filters (50/60 Hz), common average referencing (Ludwig *et al*., 2009) across all or a selected subset of channels and the ability to listen to channels of interest. These features were used to ensure that denoising was performed appropriately prior to the experiment and to determine whether data collected was electrophysiological in nature.

By default, the UI is initialized with a raster image of the sensor, producing a spatial map of activity on all channels. This enables the user to implement connectivity masks (see Methods: Array connectivity testing for more detail). Each pixel on the raster represents an individual amplifier channel, and each channel’s raw or filtered output can be viewed in a separate trace-scope, in either live-stream mode, or in threshold-trigger mode, similar to an oscilloscope.

The UI server also has an API that can be used to control the system through an external program such as MATLAB to facilitate additional analyses (e.g. channel RMS values or power spectrum plots) during recording sessions.

#### Array connectivity testing

To complete the assembly of the device, microelectrode arrays were physically pressed onto the chip while monitoring connectivity. The connectivity was assessed in the physiological saline bath by submerging electrode recording sites while applying a 2 kHz, > 2mVPP sine wave. Using the UI described previously, the pressed array was adjusted until maximal connectivity was observed and the press was stabilized with set screws on the housing within the front-end board. This completed the process of bonding the microwire electrode array to the CMOS sensor.

Next, a quantitative connectivity test was performed to determine the gain and noise of the electrically coupled array-chip device. Gain and noise were calculated using a custom MATLAB program (MATLAB 2018a, Mathworks, USA) that fetches data through the system API described previously. The gain on every pixel was calculated by applying a 2 kHz, 0.5 mV peak to peak sine wave signal in the saline bath and measuring the recorded voltage. The noise was then calculated by shorting the system reference to the saline bath (with no external signal applied). The noise measurement was band-pass filtered between 300 Hz and 6 kHz (i.e., within the spiking band). The input-referred noise was determined using the recorded root mean squared (RMS) noise values and the gain for each pixel calculated in the previous step.

The connectivity map was determined using *k*-means clustering of gain and noise values. Clusters with appropriate gain (mean > 650 V/V) and noise (mean <10 μV_RMS_) were considered optimally connected to the system, with the number of clusters set interactively by the user to achieve these thresholds. From this analysis, a connectivity mask was generated to remove unconnected channels from subsequent analysis.

### Animal Surgery

#### Rat Surgery

This study and all experimental protocols were approved by the Institutional Animal Care and Use Committee (IACUC) at the University of Texas at Austin, which follows the National Institutes of Health (NIH) guidelines for the ethical treatment of animals. Male Sprague Dawley rats (Charles River, 250-400 g) were used for implanting Argo microwire arrays. Rodents were housed in a laboratory environment on a 12-h light and dark cycle at 25°C, with *ad libitum* access to food and water.

Animals were anesthetized in an induction chamber on 3% isoflurane and then transferred to a small animal stereotaxic frame (Narishige, Japan) and maintained at 2.5% isoflurane for the duration of the procedure. Hair was removed from the head with small animal shears, Puralube (Dechra, UK) was placed on the eyes, and the surgical sites were sterilized with alternating povidone-iodine and alcohol pads prior to incision. Temperature was monitored rectally, and a heat mat was placed under the animal for the duration of the procedure with temperature maintained at around 36.0-36.5°C.

An incision was made through the scalp exposing the skull. The fascia was reflected and the periosteum was removed from the bone. A small craniotomy was made (5 × 5 mm) with a microdrill (OmniDrill 35, World Precision Instruments, USA) and the margins were removed with rat skull-appropriate bone rongeurs to extend the craniotomy to accommodate a 10 mm diameter microwire array. A durotomy was then performed with the use of microscissors. Once the durotomy was completed a micromanipulator was used to insert the Argo Microwire array into the brain and physiological saline was applied to keep the brain moist during the experiment. Argo microwires were inserted 0.7-1 mm into the brain to ensure that most of the electrodes would reside in the grey matter of the brain to enable recording of neural unit activity.

To create a reference electrode, an 80 μm diameter teflon (PTFE)-coated PtIr wire (AM Systems, USA) was de-insulated 1-2 mm from one end. The deinsulated end was then placed in the subdural space. The wire was secured in place with gel foam (Pfizer, USA) and secured on the skull with the application of UV-curable dental acrylic (Flowable Composite 101-6773, Henry Schein, USA). This limited the motion of the reference wire to reduce the possibility of picking up microphonic noise.

#### Sheep Surgery

This study and all experimental protocols were approved by the Institutional Animal Care and Use Committee (IACUC) at the Bridge PTS, San Antonio, Texas, which follows the National Institute of Health (NIH) guidelines for the ethical treatment of animals. White face Dorset Sheep (Ovis aries) that weighed 30 to 35 kg were used for this study. Food was withheld for 24 hours prior to surgery, while water was provided to the sheep *ad libitum*.

Sheep were induced by using Tiletamine/Zolazepam (Telazol, 4-6 mg/kg, IM). The sheep was intubated, and anesthesia was maintained via 1-5% isoflurane delivered in 60% oxygen and 40% medical grade air. An orogastric tube was placed to minimize or prevent ruminal bloat. Ophthalmic ointment (Paralube, Dechra, UK) was applied to prevent corneal desiccation. Thermal support was provided via a circulating warm water blanket (T/Pump, Stryker, USA) during the course of anesthesia or Bair hugger (3M, USA).

Once the sterile field was prepared, an incision was made over the skull to expose the bone and underlying fascia. The tissue was reflected, and the periosteum removed over the exposed skull. Next, a surgical microdrill (OmniDrill 35, World Precision Instruments, USA) was used to perform the craniotomy. Bone rongeurs were used to remove excess bone and fully expose the surface of the dura. A craniotomy was made (typically 3 × 3 cm) over the auditory cortex in the sheep. Typical stereotaxic coordinates were 5 mm anterior and 25-30 mm medial from bregma point for the center point of the craniotomy. After the craniotomy, a durotomy was performed to expose the pia with the use of microscissors. The surface of the brain was kept moist with the aid of physiological saline soaked gel foam (Pfizer, USA) throughout the experimental procedure.

To create a reference electrode, an 80 μm diameter teflon (PTFE)-coated PtIr wire (AM Systems, USA) was de-insulated 1-2 mm from one end. The deinsulated end was then placed in the subdural space. The wire was secured in place with gel foam (Pfizer, USA).

During the procedure, vitals were closely monitored (e.g. SPO_2_, respiratory rate). To minimize the effects of brain pulsation, end-tidal CO_2_ was typically maintained between 30-40 mmHg using mechanical ventilation. At the conclusion of the experimental procedures, animals were euthanized with an overdose of sodium pentobarbital (110 mg/kg, IV).

### In-Vivo recordings

#### Electrophysiology system

Electrophysiology recordings were acquired using the Paradromics Argo System at a sampling frequency of 32 kHz. Typically for the recording procedure, isoflurane was reduced to <2% to reduce the effect of anesthesia on neural activity.

#### Rat Action Potential Recordings

Microwire arrays between 5 mm and 10 mm in diameter (n = 5) were implanted into the somatosensory and prefrontal regions of the rat cortex. These areas were targeted for the ease of insertion of the array into the rat cortex due to the limited brain size relative to the array diameter. A successfully inserted electrode typically had a peak-to-peak noise floor amplitude of 25-30 μV.

Spiking channels of interest were evaluated in real-time with the oscilloscope and strip chart modes on the UI to confirm the presence of spike-like waveforms (typical peak width < 1 ms). We also examined specific channels of interest to observe neural firing patterns and ensure that high frequency noise was not corrupting the signal.

As a final confirmation of that the signals recorded were physiological in origin, isoflurane levels were often increased to 5% at the end of the procedure to observe the loss of neural activity and the return of the channels to the 25-30 μV noise floor.

#### Auditory Paradigms

Auditory stimuli were presented free-field through multifield speakers (MF1, Tucker-Davis Technologies, USA) approximately 10 cm from the ear contralateral to the recording hemisphere. Signals were controlled using custom MATLAB software via a data acquisition system (USB-6366, National Instruments, USA) at a sampling rate of 192 kHz, which delivered the signal through an amplifier (SA1, Tucker-Davis Technologies, USA). The speaker was calibrated using a 0.25 inch condenser microphone (PCB Piezotronics,USA).

Pink noise was presented at 80 dB Sound Pressure Level (SPL) (100 ms duration with 10 ms rise/fall cos^2^ ramp), with an inter-stimulus interval of 600 ms. For the rat, 800 stimuli were presented, and for the sheep, 500 stimuli were presented.

For the sheep, pure tones were also presented (50 ms duration with 10ms rise/fall cos^2^ ramp) at levels from 40 to 80 dB SPL in 10 dB steps, at an inter-stimulus interval of 500 ms. Pure tones ranged from 0.5 to 32 kHz with 3 steps/octave, with 20 repetitions per stimulus.

### Offline Recording Analysis

Exported HDF5 files were imported into MATLAB. Channels were bandpass filtered (using the built-in ‘filtfilt’ function, forward and backward filtering to eliminate phase delays and distortions) from 300-6000 Hz. Channels were visually inspected, and their power spectra plotted, to confirm no noise source contamination. After initial data checks, spike sorting was performed on all the channels.

Spike sorting was performed using Wave_Clus (Quiroga, Nadasdy and Ben-Shaul, 2004). Typical thresholds for neural data were set at least 3.5 times the noise threshold, calculated through Wave_Clus. In short, the noise threshold (σ) was calculated by taking the mean of the absolute value of the bandpassed signal and dividing it by 0.6745, and the spike crossing threshold was set at a minimum of 3.5σ (Quiroga, Nadasdy and Ben-Shaul, 2004). The output of the batch processing was then used to confirm the presence or absence of spike waveforms. Single units were confirmed by three metrics: (1) all neural waveforms had a peak width less than 1 ms. (2) A neural interspike interval histogram (Quiroga, Nadasdy and Ben-Shaul, 2004; Rossant *et al*., 2016) with a clear indication of a refractory period (i.e. no waveforms in the 0-3 ms bins on Wav_Clus output) was observed. (3) Clusters were clearly separated, as confirmed through the Wave_Clus user interface. Waveforms that did not match these criteria were deemed not to be single units and were not used for subsequent analysis.

#### Neural Activity Quality Metrics

To evaluate the recording quality of our isolated single units we calculated the peak-to-peak amplitude (P2P), noise, and signal-to-noise ratio (SNR). P2P was calculated by taking the peak-to-peak amplitude of median waveform of the sorted putative single unit. The noise (StdNoise) was then determined by calculating the standard deviation of the RMS noise of the band-passed signal and multiplying it by two (Kipke *et al*., 2003; Ludwig *et al*., 2011; Sohal *et al*., 2014). SNR was then calculated as P2P/StdNoise.

#### Sheep LFP recordings

##### Signal Conditioning

For LFP data analysis, all channels in the connectivity map generated immediately prior to the experiment were digitally band-pass filtered from 2 Hz to 400 Hz, with additional filtering to remove 60 Hz harmonics, and the resulting data were decimated to an effective sampling frequency of 1 kHz. To highlight evoked responses to pure tones stimuli in particular, and to further increase SNR for these recordings, the data were denoised by applying independent component analysis (ICA) (Hyvärinen and Oja, 2000). The data were first scaled and whitened using principal component analysis, before performing ICA decomposition. Next, the data were further processed by applying the Hilbert transform and evoked responses were constructed from the envelope (2-400 Hz) of the signal amplitude to account for the oscillatory nature of the LFP waveforms.

##### Pink Noise Response Analysis

Evoked responses to pink noise were compared to baseline activity for each channel. The average baseline across trials was calculated using a time window of 100 ms preceding stimulus onset to stimulus onset, and the average response was determined using a window from 20 ms to 120 ms after stimulus onset. This additional 20 ms delay after stimulus onset was to accommodate for the minimum latency that we observed for auditory responses for surface LFP recordings, similar to that previously reported in other model systems (Kajikawa and Schroeder, 2011; Escabí *et al.*, 2014; Trumpis *et al*., 2017). The ΔRMS was calculated by subtracting the RMS of the baseline window from the RMS of the signal in the response window. Channels having a ΔRMS of greater than 10 μV were considered responding channels, and evoked responses for each channel were considered statistically significantly different from baseline activity if *p*<0.05, using a multiple comparisons permutation test (Nichols and Holmes, 2002).

##### Tone Response Analysis

Responses to each tone were averaged across trials and across all levels presented, since most responses were above threshold at the lowest presented level (i.e., 40 dB SPL). The non-parametric Wilcoxon Rank Sum tests with Bonferroni correction were used to distinguish statistically significant evoked responses from baseline (*p* < 0.05/number of channels), as the data were non-gaussian. A channel was considered responsive to pure tones if it had a statistically significant response to at least two tones (i.e., 0.6 octave bandwidth), and a ΔRMS of at least 10 μV, which was considered the minimum response threshold. For these responding channels, the Best Frequency (BF) for each channel was determined as the tone which evoked the most significant response.

## Results

Here we designed and characterized the Argo system, which included a microwire electrode array of up to 31,000 channels bonded to a custom CMOS chip. Subsequent sections describe our characterization and validation of the system, which includes bench testing and performing *in vivo* experiments to record spikes in the rat cortex with 1300 microwires and surface LFP from the sheep auditory cortex with >30,000 microwires.

### Benchtop characterization of the bonded microwire-electronics system

We validated the system through bench testing physiological saline by evaluating the connectivity between the microwires electrodes and the amplifier inputs on the CMOS-sensor. Although electrodes at the periphery of the array occasionally showed lower connectivity, we were typically able to connect the majority of electrodes to an amplifier input using this process.

Connectivity tests were performed for every electrode array and sensor combination that was assembled. For 32 sensor/microwire array combinations, we obtained 71 ± 2.9% connectivity(+/-SEM) for all electrode pitches tested (60-300 μm).

The microelectrode array used for surface LFP recordings had an electrode pitch of 60 μm and the resulting 12 mm × 12 mm array contained ∼35,000 electrodes. For this array, 30,146 pixels were connected to microwire electrodes, giving 86% connectivity (Figure 5A). The gain (+/-SD) was 811 ± 21 V/V, and the corresponding input-referred noise (+/-SD) was 6.3 ± 0.5 μV_RMS_ in the 300 - 6000 Hz band (Figure 5B). From our *k*-means gain and noise analysis, 99.2% of the connected channels had < 10 μV_RMS_ band-limited input-referred noise.

**Figure 5:**
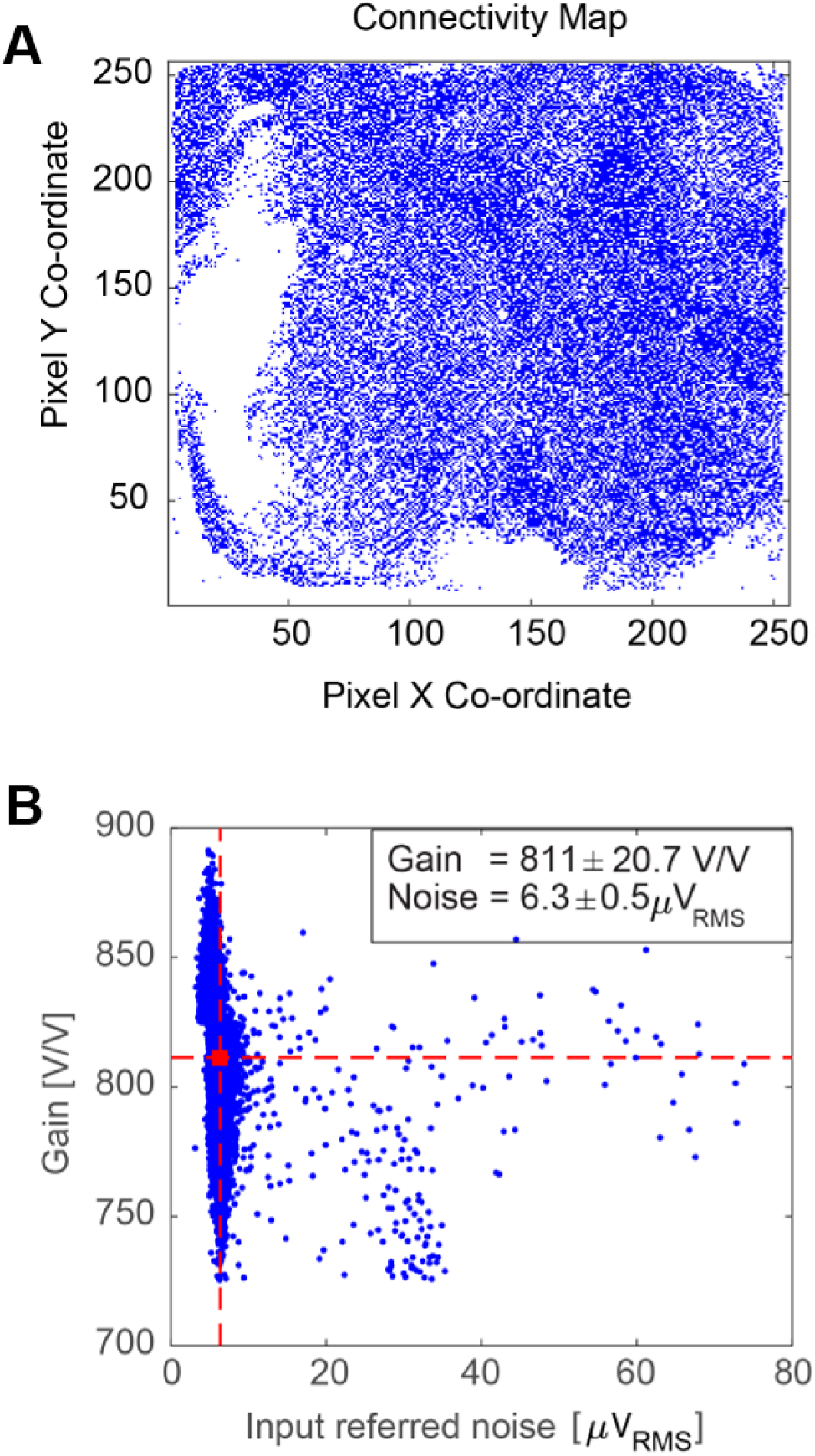
Full signal chain characterization in saline bath prior to *in vivo* experiment. (A) Raster image of the full sensor showing pixels connected to active electrodes for an array of approx. 35,000 electrodes. (B) Gain and noise distribution for the connected pixels. Channel clusters with mean gain >650V/V and mean noise <10 µV_RMS_ were selected for further analysis; here, the mean gain was 811 V/V (horizontal dashed red line), and the mean noise was 6.5 µV_RMS_ (vertical dashed red line). The final connectivity map contained 30,146 channels for analysis, resulting in a connectivity of 86%.

Across 32 CMOS sensor-bonded microwire arrays, the gain (+/-SD) was 763 ± 7 V/V, and the corresponding input-referred noise (+/-SD) was 7.5 ± 0.4 μV_RMS_ in the 300 - 6000 Hz band. Thus, a large array of functional electrodes can be created using this CMOS sensor-microwire bonding approach with high connectivity across the sensor and good gain/noise characteristics for connected channels.

### Rat Cortical Recordings

To validate our system *in vivo*, we show data from a 1300 microwire array (10 mm array diameter, 18 µm wire diameter, 200 µm spacing, 1 mm length), the largest microwire array implanted into the rodent cortex to date. The array was implanted into the somatosensory and prefrontal regions of the rat cortex for ease of insertion of the large array due to these areas being on a flatter part of the rat cortex and requiring minimal rotational manipulation of the headstage for insertion. Together, this allowed for successful array insertion normal to the surface of the brain. Using this array, we were able to isolate 791 single units (Figure 6A,C). Mean +/-SEM peak-to-peak amplitude, noise and SNR values obtained across all units were 130 µV +/-59.7, 10 µV +/-2.2, 8.9 +/-1.9 respectively (Supplementary Figure S1). Therefore, high fidelity neural activity can be recorded with the Argo recording system and associated microwire arrays along with a high unit count.

**Figure 6.**
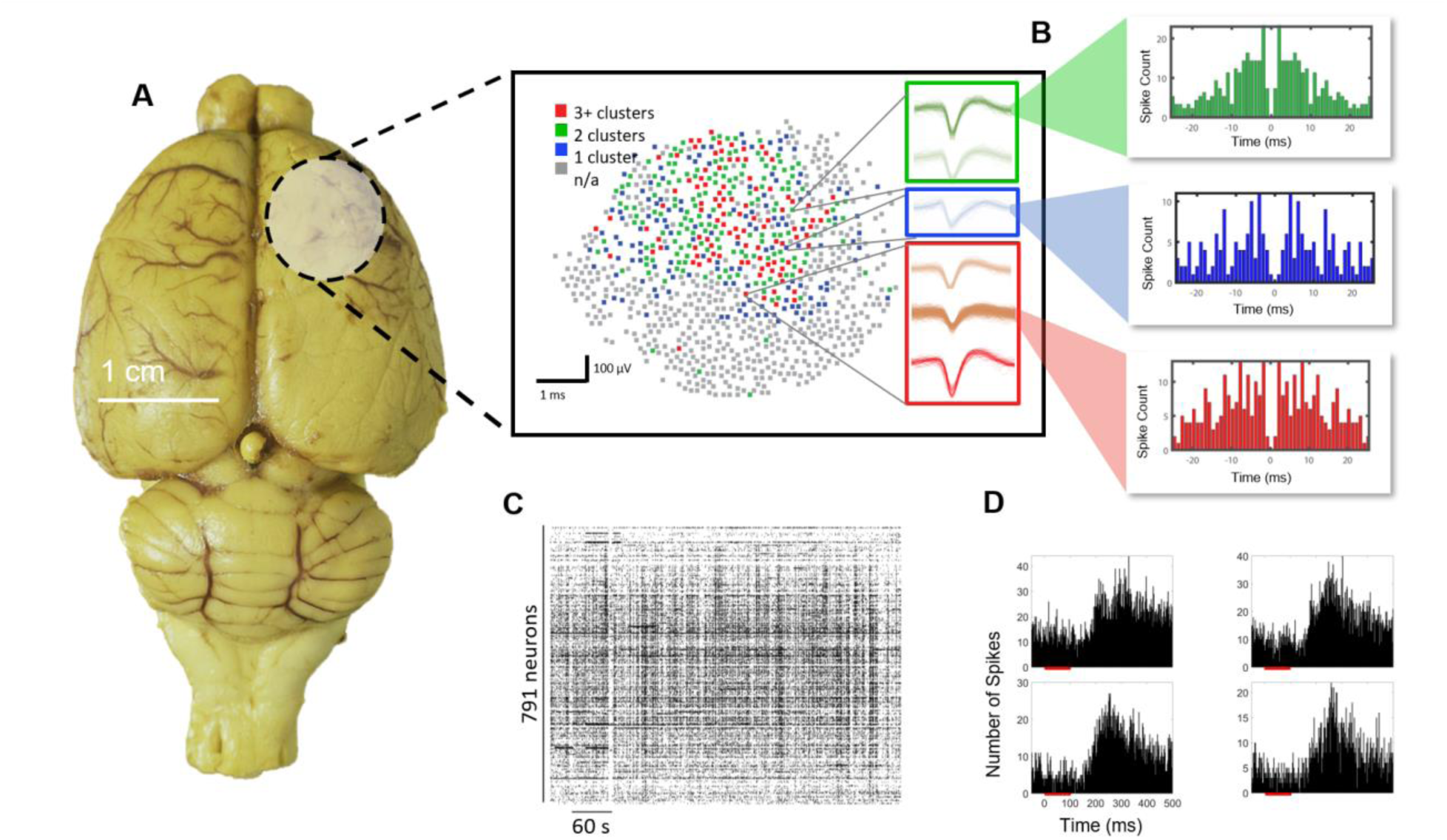
Rat cortex spike recordings. (A) Distribution of units found over the implantation of 1300 microwires in the rat cortex. The circular shaded region shows the location of the craniotomy and the brain region that was implanted, and the map shows channels with 0-3+ spike clusters. (B) Example autocorrelogram from specific highlighted units, showing the emergence of well isolated single units. (C) A total of 791 single units as shown by the spike raster were found in the recording with good temporal dynamics associated with neuronal firing under anesthesia. (D) Example post-stimulus time histograms responses of single units in the somatosensory cortex to auditory pink noise stimuli (red bar). A latency of around 200 ms is observed, as expected from somatosensory neurons responding to auditory stimuli (Maruyama and Komai, 2018).

In addition to baseline recordings of spontaneous activity, pink noise auditory stimuli were presented to evoke somatosensory responses. Auditory stimuli were chosen in lieu of other sensory stimuli to avoid artifacts typically introduced by more direct somatosensory stimulation paradigms such as physical manipulation of the animal. We observed single units from somatosensory cortex that responded to auditory stimuli at an expected latency of 200 ms (Maruyama and Komai, 2018), further confirming neural activity (Figure 6D).

The rat cortex typically does not favor insertion of high density electrodes due to the brain size and high possibility of vascular insult (He *et al.*, no date; Kozai *et al*., 2010; Blinder *et al*., 2013) resulting in damage to neural tissue. However, we were able to obtain high fidelity recording with high unit count (Figure 6A,B) owing to our microwire design (electro-sharpening) and ease of insertion into the rat cortex.

### Sheep Surface Recordings

While the rat cortex was ideal to validate the Argo system’s ability to record neural action potentials, the size of the microwire array was limited by the small size of the rodent brain. As a result, recordings would always be limited to 1000-2000 electrodes, substantially smaller than the maximum channel count of the Argo system. We therefore developed microwire arrays for surface LFP recordings with over 30,000 connected channels in order to evaluate surface potentials from sheep cortex (n = 2). The ∼30,000 channel limit was the result of a tradeoff between electrode spacing, ideal microwire diameter and recording site area to allow for the acquisition of surface LFP at an appropriate spatial resolution.

The microwire arrays used in these recordings had a pitch of 60 µm, with an overall array size of 12 mm × 12 mm. We targeted the array over the auditory cortex (centering at 25 mm lateral and 5 mm anterior of bregma). LFPs were recorded at the surface of the cortex in response to pink noise (Figure 7) as well as pure tones (Figure 8). To quantify differences in evoked responses to pink noise compared to baseline activity, we first calculated the ΔRMS for each channel in the 2 Hz to 400 Hz band (Figure 8A). As an initial measure of activation, we considered channels to be responsive if the evoked activity was at least 10 µV RMS higher than baseline, which is twice the determined noise floor. In one sheep, 21,963 channels of the 30,146 total channels (73%) showed an increase in power above threshold during the pink noise stimulus, with 16,412 (54%) of those channels’ reponses being statistically significant. (Multiple comparisons nonparametric permutation test, *p* < 0.05) (Nichols and Holmes, 2002). In a second sheep, we found that 17,233 of 31,239 total channels (55%) showed increased power, with 12,590 (40%) channels having a statistically significant response.

**Figure 7.**
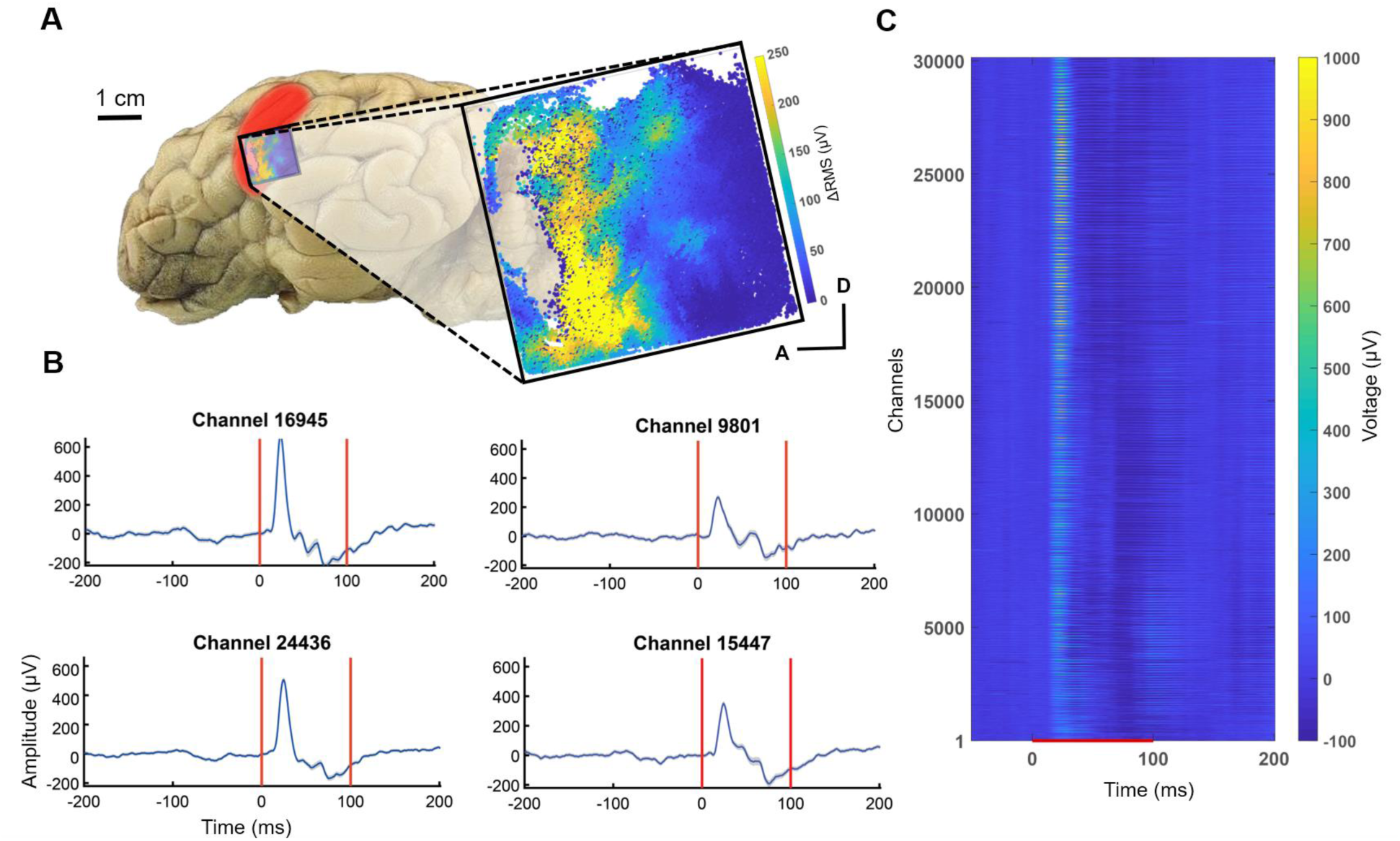
Pink noise-evoked potentials across the 30,000-channel microwire array from the sheep auditory cortex (red shaded area, described in (Gierthmuehlen *et al*., 2014)). (A) Location of the array (blue shaded boundary) for the auditory cortex surface recording and a map of ΔRMS (i.e., response RMS minus baseline RMS) for individual channels in response to pink noise. With the ΔRMS of at least 10 µV, 21,963 of these channels were responders to pink noise, A (anterior) and D (dorsal) for the channel location in the array. (B) Evoked potentials (mean +/-SEM) from representative responding electrodes. (C) Trial-averaged evoked potentials of all 30,146 channels in response to pink noise. The red line denotes when the stimulus was presented. Channels were ordered spatially by the pixel location.

**Figure 8.**
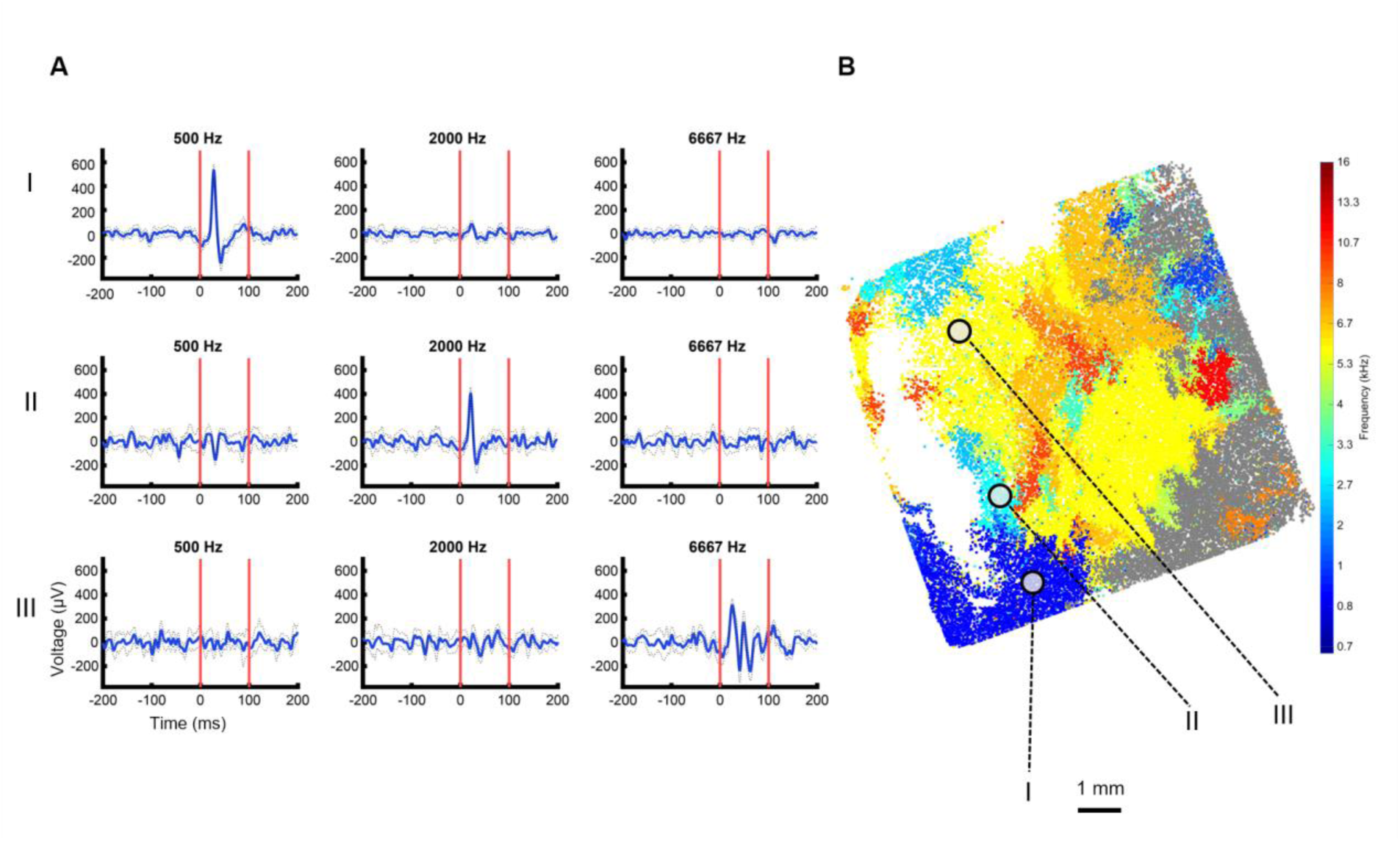
Frequency-specific responses with emerging tonotopic organization for the sheep auditory cortex. (A) Evoked potentials from electrodes across the microwire array demarcated from (I-III) are characteristic of frequency specific responses. (B) Array map with electrodes color coded for BF. Non-significantly responding electrodes shown in gray (Wilcoxon rank-sum, *p* < 0.05, Bonferroni corrected).

When presenting pure tone stimuli, we also observed frequency-specific responses (Figure 8), where 22,206 of 31,239 (71%) channels had at least 10 µV RMS activity above baseline and were significantly responsive to at least 2 tones (i.e., a minimum bandwidth of 0.6 octaves) (p < 0.05 Wilcoxon Rank Sum test, with Bonferroni correction). We then determined the BF for each channel by ranking the level of response to each frequency played. The BF for each channel was used to estimate the tonotopic organization across the cortex (Figure 8B). This was replicated in another experiment, where 26,054 of 30,146 (86%) channels were responsive, with a similar tonotopy across the cortex (Supplementary Figure S2).

## Discussion

Here we have described and characterized a neural recording system based on microwire electrodes compressively and reversibly bonded to a custom-designed CMOS amplifier array. We have demonstrated that these recording devices provide large gain and a low noise floor and can scale to a very large number of recording sites with optimal gain and noise distributions. This makes such devices suitable for massively parallel electrical recordings of spikes and LFP in animal cortex.

To validate this recording capability, we have demonstrated the largest microwire electrode array-based recordings in both rat and large animal cortex to date. We show the ability to record from over 30,000 channels simultaneously at full acquisition rates (32 kHz) in the sheep auditory cortex with good responses to auditory stimuli. Further, we show the ability to record high fidelity spikes in the rat cortex with our microwire based technology. Both results highlight and validate the capability to simultaneously record large channel count neural data at acquisition rates of 32 kHz with the Argo system.

### Sheep Neural Recordings Demonstrate 30,000 channels of simultaneous acquisition

From the sheep cortex, we recorded surface LFP on over 30,000 channels with a 12 mm × 12 mm array, which is the largest neural recording in a large animal to date. Other studies from µECoG in large animals and humans range from 16-294 channels (Chiang *et al*., 2020), therefore our system provides an order of magnitude increase in channel count. We performed surface LFP recordings using a high-density electrode array to mitigate the risk of damage to neurons that could have occurred upon insertion of a high-density penetrating array, resulting in poor recording quality.

To the best of our knowledge, this is also the first *in vivo* demonstration of evoked responses to auditory stimuli in sheep. The auditory cortex areas identified here were similar to those previously reported with histological tracing experiments in sheep (Michaloudi *et al*., 1986; Gierthmuehlen *et al*., 2014), which reported the auditory cortex to be approximately 1 cm × 2 cm. Responses to auditory stimuli typically had a latency of approximately 20 ms after stimulus onset, which is expected with surface LFP recordings from other relevant animal models (Kajikawa and Schroeder, 2011; Escabí *et al*., 2014; Trumpis *et al*., 2017). We did observe slightly different responses to tones across the microwire array between experiments, with one experiment revealing more channels with higher BFs. This is likely due to small differences in placement, especially since the array could span approximately half of the auditory cortex, and individual animal variation. Moreover, this tonotopy and sound-evoked responses were evident despite the acoustically noisy environment (i.e., an acute operating room setting rather than the traditional sound booth environment used in traditional auditory experiments), which could have led to increased variability in responses. However, even with this variation, we were able to find emerging tonotopy in the sheep auditory cortex.

### Rat Neural Recordings Reveal High-fidelity Action Potentials

We demonstrate the ability to record from 791 single units in the rat cortex. This is significantly more than other microwire technology that has been used to record from the rodent cortex, where the number of neurons ranges from 20-240 units in an acute setting (Guitchounts *et al*., 2013; Pfeiffer and Foster, 2013; Chung *et al*., 2019; Massey *et al*., 2019; Obaid *et al*., 2020, no date). We found the mean SNR across all channels to be close to 9, indicative of high fidelity recordings given that typical values using this metric define good SNR between 3 and 6 (Ward *et al*., 2009; Ludwig *et al.*, 2011; Kozai *et al*., 2012; Sohal *et al.*, 2014; Xu *et al*., 2018). Here we demonstrate a system that is capable of producing low noise and high captured signal amplitude from single units. This is enabled in large part by the combination of tunable high pass and low pass filters that allow us to set the input bandwidth to reject both low and high frequency noise and remove aliasing effects.

Furthermore, the design of the microwires enables good performance for neural recordings. Specifically, our microwire electrodes have tip based recording sites, which are known to produce better recording quality compared to those sites that are along the length of the shaft, like some multi-depth silicon probes (Kipke *et al*., 2003; Jun *et al*., 2017). The electrosharpening of the tips also allows for ease of high-density array insertion into the brain (Obaid *et al*., 2020, no date) as has been demonstrated by our neural recordings, where we can successfully insert arrays into the rat cortex.

### Microwire-CMOS devices as Next Generation Brain-Computer Interfaces

Microwire-CMOS devices combine the robustness and longevity of traditional microwire electrodes with the advantages of active CMOS probes. Specifically, the design overcomes three of the greatest drawbacks of using microwire electrode arrays for neural recording. 1) By amplifying close to the signal source, the design reduces both input capacitance and noise pickup. 2) By providing on-chip multiplexing, the design allows for a much smaller number of lead wires and connectors than electrode sites. 3) Bonding pre-assembled arrays of microwires onto a CMOS sensor array provides a much simpler method of connecting the microwires than the traditional hand-wiring approach.

Here we demonstrate that microwire-CMOS is indeed scalable to tens of thousands of channels, but the current system is limited to acute, head-fixed preparations due to the size of the electronic components that lie downstream of the CMOS amplifier array. The clear next step is to develop a new ASIC and downstream architecture that are compatible with a floating microwire array configuration. Such a device would have a form factor similar to the Utah array but allow for greater electrode density, reduced tissue damage and insertion forces owing to smaller electrode diameter (Patel *et al*., 2015; Obaid *et al*., 2020, no date), and reduced interface cable thickness due to on-chip multiplexing of channels. Development of such a device is already underway.

## Acknowledgments

This work was funded by a Small Business Innovative Research grant from the National Institute of Health Brain Initiative (5R43MH110287-02) and by the Defense Advanced Research Projects Agency’s Neural Engineering System Design program (N66001-17-C-4005). The authors would like to thank Prof. Andreas Schaefer for feedback and discussions, and Dr. Marike Zwienenberg, Linda Talken, and Amy Lesneski (UC Davis) for help in developing sheep surgical protocols. **Competing interests:** K.S., A.A.K., A.P.S., T.M.S, Y.N., A.T., P.O., C.L., D.P., K.N., K.M.B., S.S., M.S.H., B.K., M-E.S.H., R.J.E., I.M., D.F., A.M.S., V.G., Y.K., M.S., H.S.S., M.R.A. are current or former compensated employees or consultants of Paradromics, Inc., a brain-computer interface company. P.G., A.B-F, S.V., A.V.K, B.L., B.D. are compensated employees or consultants of Caeleste, CVBA, a circuit design company.

