## Supplementary Material for "The Argo: A 65,536 channel recording system for high density neural recording *in vivo*"

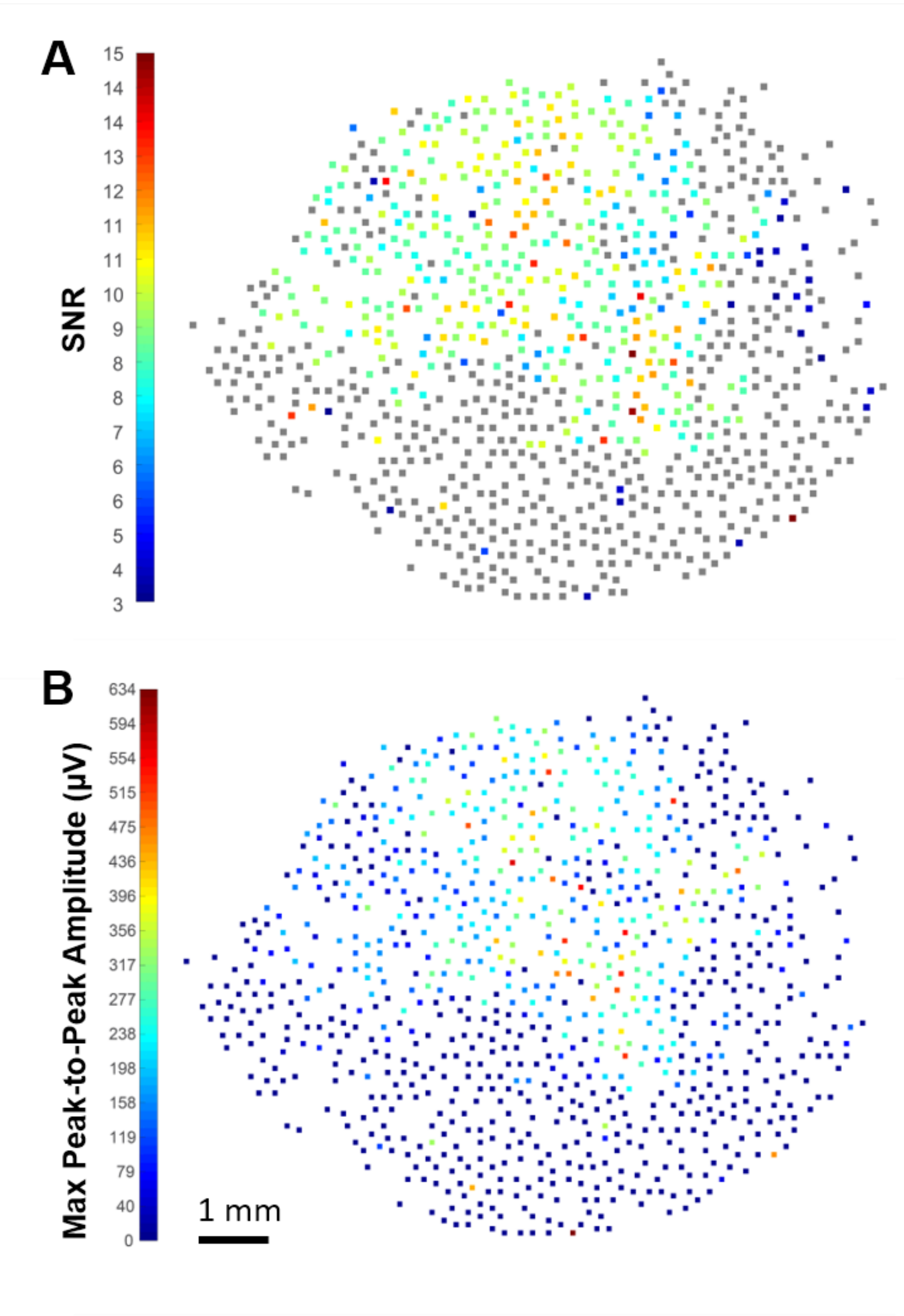


**Figure S1**. Distribution of both SNR (A) and Max Peak to Peak amplitude (B) for our 1300 microwire recording in the rat cortex. High SNR and P2P amplitudes can be observed indicating the presence of high-fidelity neural recordings.

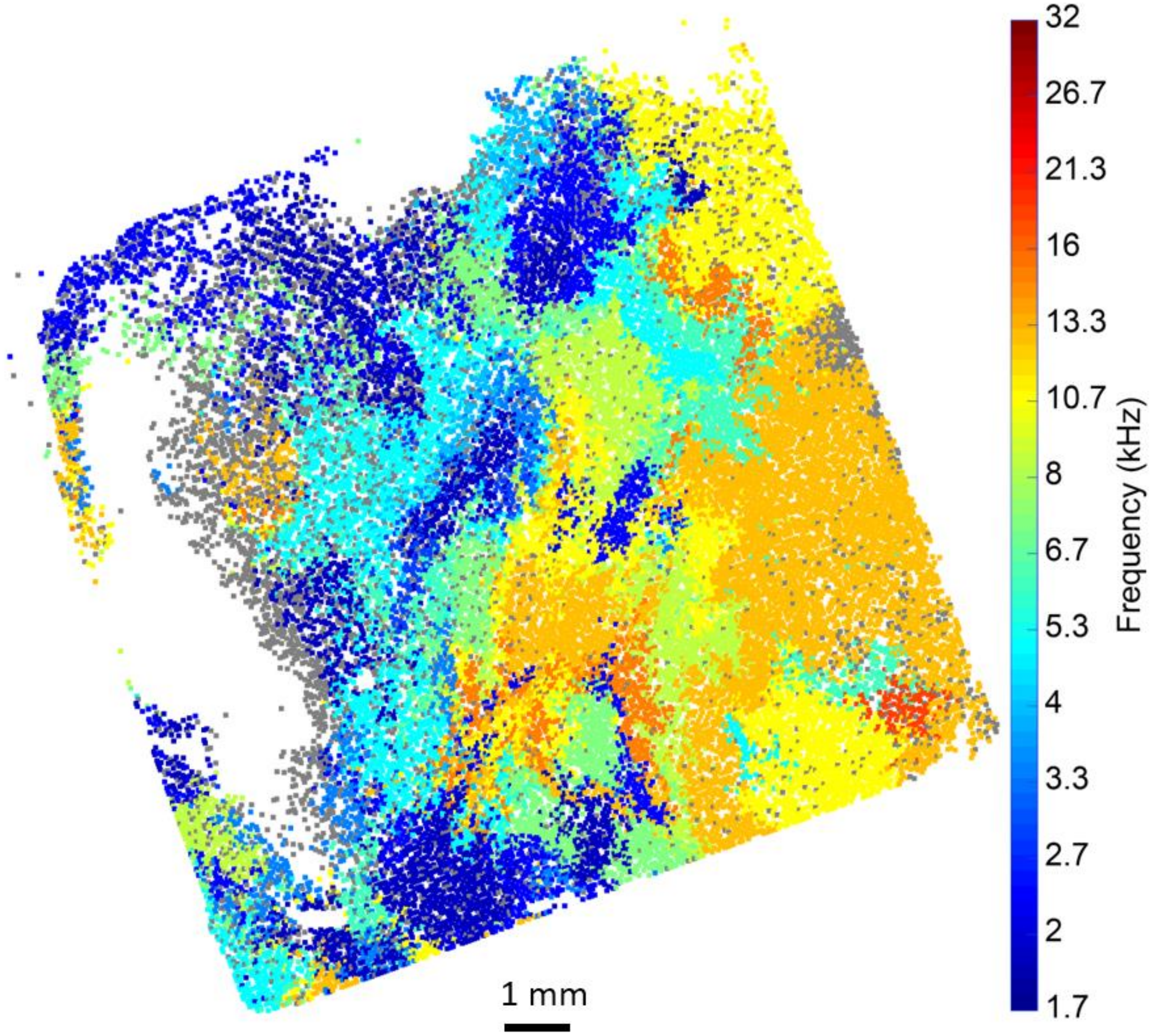


**Figure S2** Array map with electrodes color coded for Best Frequency (BF). Non-significantly responding electrodes shown in gray (Wilcoxon rank-sum, $p < 0.05$, Bonferroni corrected).

| Parameter | Value |
|---|---|
| Power Mode | AC |
| Slew Rate | 60 V/ms |
| Current Limit | 10 A |
| Overcurrent Delay | 15 s |

**Table S1: Electrosharpening system parameters.** The Chroma system was chosen for its high current limit of 10 A.

| Step | Process Start Distance Below Surface (mm) | Voltage (V) | Frequency (Hz) | Process notes |
|---|---|---|---|---|
| 1 | 0.7 | 35 | 60 | Time: 8s |
| 2 | 1.2192 | 35 | 60 | Time: 10s |
| 3 | 1.2192 | 30 | 60 | Velocity: 0.07 mm/s |
| 4 | 1 | 5.7 | 51 | 5 pulses, 0.5 s long, 20% duty cycle<br>No wire movement |

**Table S2: Microwire sharpening process parameters.** In each of the first three steps, the wire tips start below the surface, and are drawn out of solution to preferentially remove material from the tip. The first step is used to ensure the wires have equal length to ensure consistency in subsequent steps. The second step is a coarse electrosharpening step in which material is removed rapidly, and the third step is a fine electrosharpening step, in which the voltage and stepper motor speed are reduced to ensure the tapers are consistent and smooth. In the fourth and final step, the wire tips are held at 1 mm below the solution surface, and five pulses are provided to polish the electrode tips. In the first and fourth steps, the z-position is set to zero at the solution surface using a resistance measurement. All distances are relative to this surface. Between each step, the peristaltic pump is run at 50 rpm to evacuate reaction products.
